# Automated 3-D mapping of single neurons in the standard brain atlas using single brain slices

**DOI:** 10.1101/373134

**Authors:** Jun Ho Song, You-Hyang Song, Jae-Hyun Kim, Woochul Choi, Seung-Hee Lee, Se-Bum Paik

## Abstract

Recent breakthroughs in neuroanatomical tracing methods have helped unravel complicated neural connectivity in whole brain tissue at a single cellular resolution. However, analysis of brain images remains dependent on highly subjective manual processing. In the present study, we introduce AMaSiNe, a novel software for automated mapping of single neurons in the standard mouse brain atlas. The AMaSiNe automatically calibrates alignment angles of each brain slice to match the Allen Reference Atlas (ARA), locates labeled neurons from multiple brain samples in a common brain space, and achieves a standardized 3D-rendered brain. Due to the high fidelity and reliability of AMaSiNe, the retinotopic structures of neural projections to the primary visual cortex (VISp) were determined from single and dual injections of the rabies virus onto different visual areas. Our results demonstrate that distinct retinotopic organization of bottom-up and top-down projections could be precisely mapped using AMaSiNe.

## Introduction

In the last few decades, research in systems neuroscience has increased and benefited from the advances in molecular and genetic engineering techniques. Using various purpose-specific techniques, neurons can be labeled and traced in both anterograde/retrograde directions ^1–4^. Subcellular neuronal components, such as ion channels expressed in specific types of neurons ^5–7^ and connectivity-specific synapses ^8–11^, can be readily manipulated. Emergence of powerful tracing techniques has enabled us to disentangle highly complicated neural circuits of diverse brain regions ^12–17^.

Despite such breakthroughs in single-neuron level tracing techniques, the analysis of obtained images still relies, in general, on old methods such as manual counting of the number of labeled neurons. Specifically, conventional experiments of mouse brain image analysis typically begin with finding a brain atlas image that is visually similar to the experimentally obtained image **(Fig. 1a)**. Then, the region-of-interest (ROI) boundary is manually drawn and the number of labeled neurons in ROIs is counted. However, this conventional analysis has several critical issues. First, manually selected ROI boundaries are highly subjective and susceptible to human error. This issue can arise from many factors that cause distortion in sectional views of brain samples that have undergone histological processes, including imperfect slicing angles that may vary across samples **(Fig. 1b)**. In addition, the commonly used 2-D reference atlases (e.g. ref.18) are affected by this slicing angle issue, thus human intervention, i.e. experimenters’ subjective decisions on finding corresponding slices in the atlas, has to resolve the discrepancy between the atlases and obtained slice images to match them. These subjective judgements inevitably lead to erroneous analysis results, especially in mice brains that lack any clear landmark structures, such as sulci and gyri.

**Figure 1.**
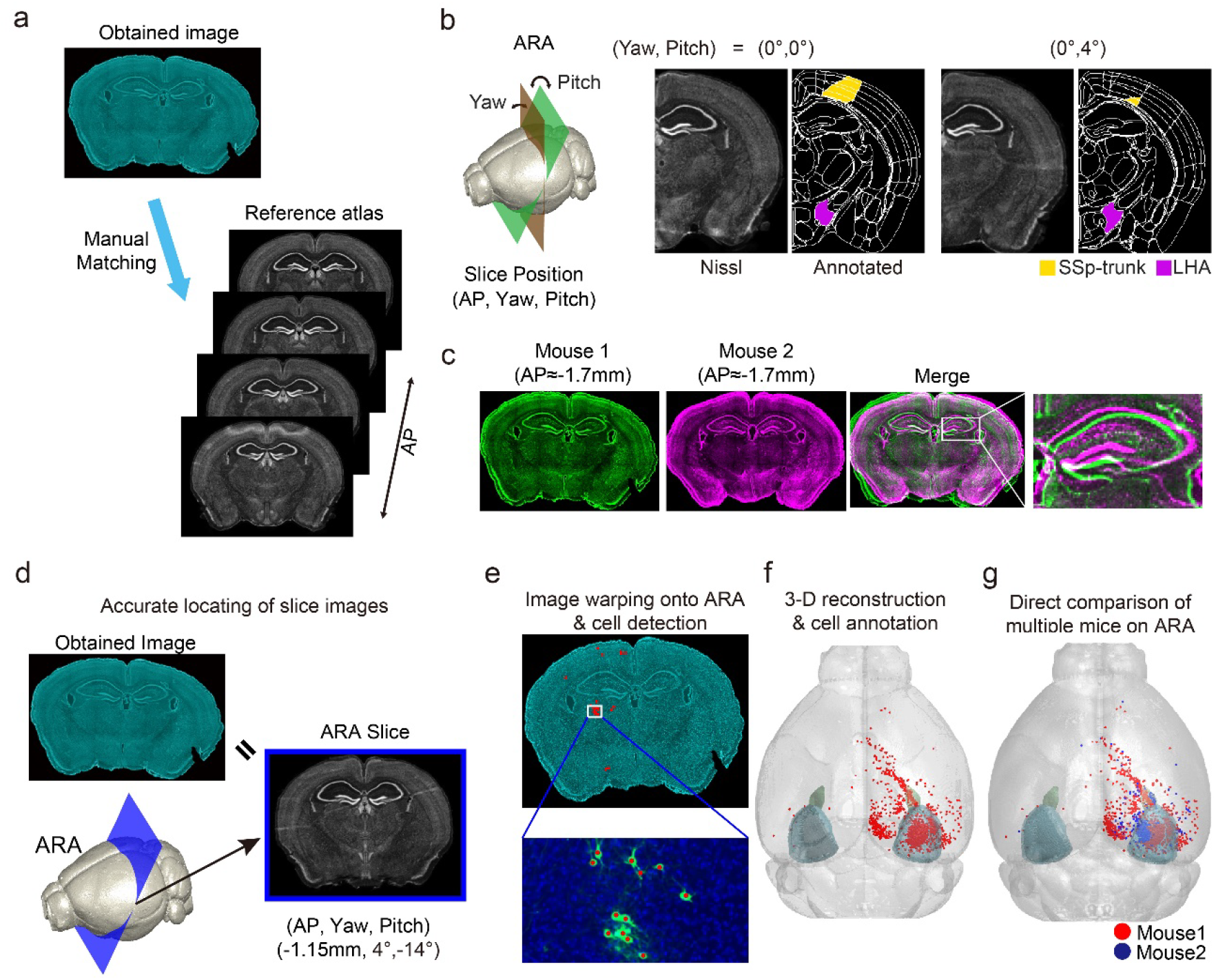
Design of AMaSiNe to overcome the limits of conventional mouse brain image analysis. (a) Conventional analysis of brain images. Investigators visually compare images obtained from experiments (top) to 2-D reference atlas slices (bottom) to annotate ROIs. (b) Inaccuracy due to variations of slicing conditions such as yaw and pitch angles, and AP position (left). Neglecting any of the position parameters results in inaccurate estimation of ROIs (middle and right). Sample images prepared from virtually slicing ARA at different tilt angles are similar in global view (Nissl images), but each annotated region differs noticeably (annotated images; SSp-trunk for primary somatosensory area for trunk; LHA-lateral hypothalamic area). (c) Practical issues regarding direct comparison of different data sets. Superimposition of two slices from different brains at similar AP positions (green and magenta) shows a mismatch (merge). (d–g) Design of AMaSiNe for accurate brain slice analysis. (d) AMaSiNe compares obtained images to virtual slices of ARA and finds their matching position in ARA. (e) Original images are warped onto the corresponding ARA space and the position of single neurons is localized. (f) Calibrated location of neurons is projected onto the 3-D reference ARA space and annotated to different ROIs. (g) Direct comparison of neural distribution from different brains using AMaSiNe.

Second, even if the ROI boundaries are set flawlessly, simply counting the number of labeled neurons in each slice image provides only the information regarding the 2-D distribution of neurons in the slicing plane. Consequently, much information regarding the complete 3-D organization of neurons is lost. To measure the neuronal distribution in non-slicing axes, reconstruction of the distribution of labeled neurons on a standardized 3-D brain atlas is required, however this is not readily achieved due to deformations of brain sections during tissue preparation. Furthermore, the sizes and handling conditions of individual brains such as slicing angles and structural deformation vary, thus, their neuronal distributions in 3-D cannot be directly compared **(Fig. 1c)**. Lastly, these manual analysis processes of brain images are highly laborious, thus, studying neuronal organization on a whole-brain scale is a formidable task.

Diverse approaches have been suggested to address these issues by implementing new hardware or image processing methods ^13, 19–25^. For example, Ragan and colleagues devised the serial 2-photon tomography ^21^, which aims to minimize tissue deformation during preparation by combining a microtome with a 2-photon microscope. Then, image data from multiple brains are warped onto a common 3-D space allowing comparison of different brains. However, the warping process is performed on a volume-to-volume basis, requiring two whole brains reconstructed in 3-D. Therefore, brains must be sliced from their anterior start to the posterior end, regardless of how small the compared region is.

In another recent study, an interactive framework that obtained brain images analyzed on a slice-to-slice basis with a scale-invariant atlas was introduced ^22^. However, the software required a manual choice of the atlas slice that best matches the observed image and does not account for the errors in the slicing angles. Therefore, without these existing issues resolved, current approaches are inevitably susceptible to common human errors.

To solve these problems, we developed a novel analysis algorithm for automated 3-D mapping of single neurons (AMaSiNe) in the standard brain atlas using single brain slices of mice. Using the Allen Reference Atlas (ARA) ^26^, AMaSiNe finds the alignment condition of obtained brain images including slicing angles and location in the anterioposterior (AP) axis. Then the software reconstructs single neuron positions in a 3-D reference space from the calibrated brain slices, automatically sets the ROIs of each brain area, and counts labeled neurons in each ROI **(Fig. 1d-e)**. From the image-to-image matching scheme, our tool can achieve desired accuracy even when very few images are available. Furthermore, data sets from different mice are successfully compared on a common 3-D space because the reconstruction is performed with the ARA **(Fig. 1f-g)**. We further illustrate that AMaSiNe can delineate the 3-D spatial organization of neurons that project to the primary visual area (VISp) in various ROIs, such as the dorsal lateral geniculate nucleus (LGd) and other higher cortical areas in the cortex.

## Results

The AMaSiNe software operates as follows: Given a set of mouse brain slice images, the software estimates the parameters for alignment condition of each slice image, such as the slicing position in the AP axis and yaw and pitch angles. The obtained image is warped onto the cross-sectional image of the Allen Reference Atlas (ARA) ^26^ that corresponds to the estimated slicing condition and calibrated to match the ARA reference image. From the warped image, neurons are automatically detected and labeled. Finally, the spatial organization of the labeled neurons is reconstructed in 3-D ARA and cells are annotated in the corresponding ROIs. Because the 3-D reconstructed brain precisely matches the ARA, AMaSiNe allows studying neural circuits in a standard space without the need to slice the whole brain. To achieve this goal, AMaSiNe is composed of a series of processing modules (**Fig. 1d–g**; **Fig. S1** shows the algorithm pipeline in detail).

### Localizing Brain Slice Images

One of the major sources of error in the manual analysis of brain slice images is the visual matching process between experimentally obtained images and 2-D atlas images (see ref. 18). Unlike standard atlas images produced under a specific condition, experimental conditions for tissue preparation vary trial by trial. For example, the alignment condition of each slice image, such as the slicing position in the AP axis and yaw and pitch angles, cannot be identical in each experiment. Inconsistencies in experimental conditions inevitably require human decisions during analysis, rendering the analysis results more vulnerable to individual interpretation.

Therefore, in the first stage of our software analysis, the three parameters of the brain slice images had to be estimated: yaw and pitch angles of the section and the location of the section along the anterior-to-posterior (AP) axis of the brain. Searching these parameters was performed by comparing an image from the experiment with multiple cross-sectional ARA images sliced at various AP positions and angles **(Fig. 2a)**. Once the ARA slice most similar to the original image was found, the position parameters of the ARA slice was designated as that of the obtained image. To perform this process, image similarity was quantified by matching the local features between ARA slices and the original image. After preprocessing the contrast adjustment, edge-sharpening, and downsampling to 50 μm, each slice image was linearly transformed to roughly match the target ARA slice **(Fig. 2b**; see **Methods** for detailed procedure). Next, structural features of the local area were extracted from the obtained image and ARA slice images by the speeded-up-robust-features (SURF) algorithm^27^ and grid-point sampling at the interval of 200 μm, respectively **(Fig. 2c)**. The purpose of grid-sampling in an obtained image was to ensure evenly spaced feature-matching and avoid excessive local weighting during similarity quantification. Finally, similar features from both images were matched and the image similarity score (ISS) computed with the histogram-of-oriented-gradients (HOG) vectors ^28^ **(Fig. 2d**; see **Methods** for the feature-matching procedure and mathematical definition of ISS). From the estimation of ISS between the original image and multiple ARA images sliced using different position parameters, the location parameters such as yaw and pitch angles and AP position of the ARA slice with the highest ISS were set as the position of the original image **(Fig. 2e)**.

**Figure 2.**
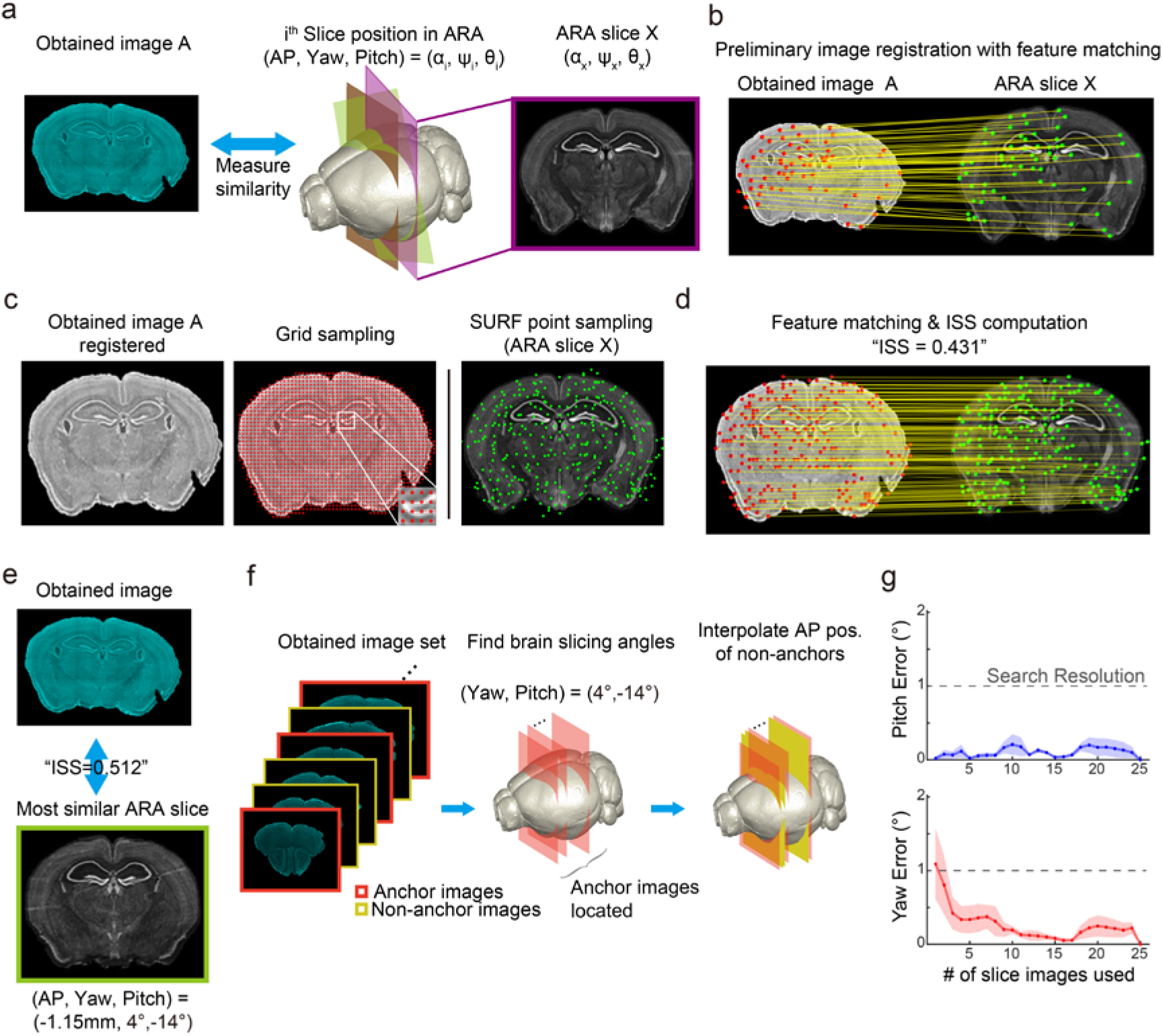
Localization of brain slice images onto the standard brain atlas (ARA) (a) Process for slice localization. AMaSiNe compares the similarity between original image and ARA images at various positions. (b–d) Calculation of image similarity score (ISS). Original image is downsampled as 50 μm/pixel resolution and edge-boosted, then linearly warped onto the ARA slice for comparison (b and c left panel). Feature points are extracted by grid sampling at a 200 μm interval in the original image and SURF detection in ARA slice, respectively (c middle and right panel). (d) Feature points in the image pair are matched based on the similarity between HOG descriptor of each feature point and ISS is calculated. Note that matched feature points in the right hemisphere are missing due to the structural dissimilarity of the hippocampi in the two images. (e) Choice of ARA slice most similar to the original image. AMaSiNe computes ISS of multiple ARA slices and finds the best matching slice that gives the highest ISS. (f) Estimation of slicing angles in an individual brain. Slicing angles are identical or very similar within a brain, thus, they can be found from a few sample images (anchor images) with less computational cost (left and middle). Non-sampled images are then interpolated from anchor images (right). (g) Accuracy of AMaSiNe for finding slicing angles. Different number of slice images were used for estimation and their errors calculated (4 brains, 25 images sampled per brain; mean ± SEM; see also Supplementary Fig. 2).

The advantage of our feature-based similarity comparison is that all local structural aspects of a brain slice are considered to compute the precise comparison of slice images, unlike individual interpretation based on visual comparison of some salient features. Furthermore, with this strategy, AMaSiNe can find the accurate parameter of slice alignment, even when a part of the image is lost or damaged during tissue preparation. Using the feature-based image-to-image comparison method, we confirmed that AMaSiNe produces consistent results with slice images stained utilizing various methods, such as the most common, DAPI, Nissl, and autofluorescence **(Fig. S2)**.

To improve computational efficiency, we implemented the following strategy: First, the three-step searching, a popular derivation of the greedy search algorithm, was applied to find the most similar ARA slice (**Fig. 2e**). Next, for the initial estimation of alignment parameters, obtained images and ARA were downsampled to 50 μm × 50 μm resolution. We confirmed this resolution was acceptable because higher resolution did not improve the accuracy of results (data not shown). Lastly, instead of searching parameters of all obtained images, AMaSiNe allows sampling of slice images from a single brain at the interval of approximately 250 μm – 1000 μm and interpolates the parameters of unsampled images from those sampled **(Fig. 2f)**. The validity of this approach was confirmed from the test for determining the minimum number of sample images to find the correct slicing angles **(Fig. 2g)**. We observed that any two images from different positions were sufficient to reliably find the slicing angles with the errors of estimated alignment angle less than 1° on average using our tissue preparation setting.

### 3-D Reconstruction of the Brain and Cell Annotation

After obtaining the alignment parameters of original slice images in ARA, AMaSiNe warped the images onto its corresponding ARA slice and compensated for the structural deformation during tissue preparation **(Fig. 3a)**. The warping was based on the feature-to-feature matching in four steps: a similarity transformation (linear), two affine transformations (linear), and a local-weighted-mean transformation (non-linear) ^29^. The transformation parameters, three matrices for linear transformations and the locations of matched features for non-linear transformation, were computed from the matched SURF points in ARA slice and obtained images. The obtained images at their original pixel resolution were geometrically transformed with these parameters. After aligning the warped images, the original slice images were successfully transformed to reconstruct 3-D positions of the neurons in the standard ARA space **(Fig. 3b)**.

**Figure 3.**
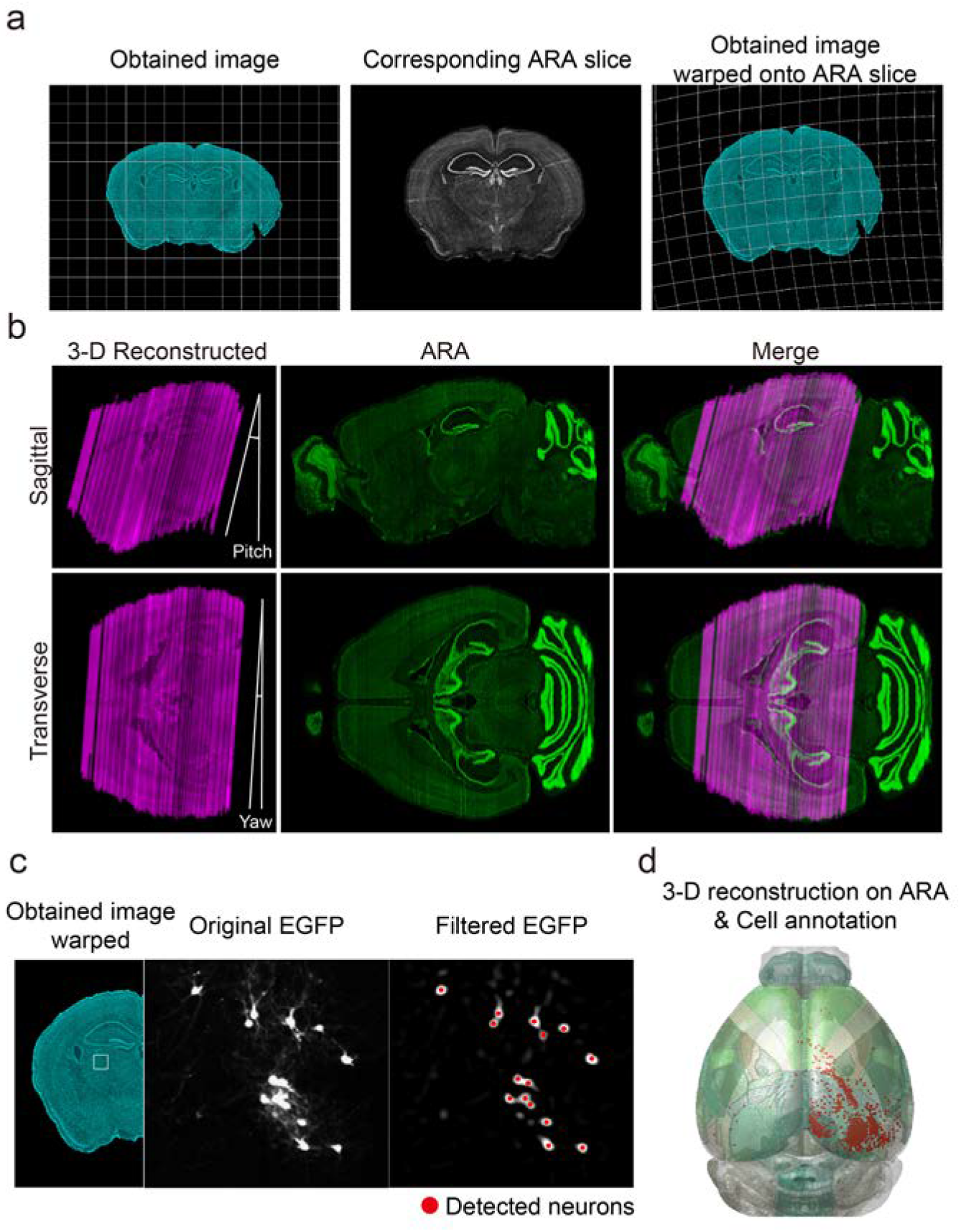
3-D reconstruction of single neurons in the brain. (a) Registration of experimentally obtained images. Original images are warped onto their corresponding ARA slice on a feature-to-feature basis. (b) Alignment of slices for 3-D reconstruction of the brain. Because individual images are registered to match their corresponding ARA slice, the reconstructed brain accurately fits into the 3-D ARA space. (c) Labeled neurons are automatically detected from the warped images and their positions recorded. (d) Annotation of located neurons. Detected neurons can be automatically allocated to each region of the reconstructed 3-D brain in the ARA space.

Once the image registration process was completed, the location of the labeled neurons’ somas in transformed images were measured **(Fig. 3c)**. A series of filters, including Fermi and difference-of-Gaussian filters, were first applied to enhance the soma’s edge. Then, circular structures, namely the labeled somas, were located using Hough transformation ^30, 31^. AMaSiNe allows users to tune the parameters for cell detection, such as the diameter range of labeled somas, for flexible application of the algorithm for soma and/or neuropil detection ^32, 33^. Because the 3-D reconstructed brain accurately fit into ARA, detected neurons in each slice were directly positioned on ARA, the common 3-D reference space. Utilizing this information, AMaSiNe allocated labeled neurons into their corresponding ROIs using the 3-D annotated ARA **(Fig. 3d)**.

### Topographic Organization of Inputs to Visual Cortex

Orderly architecture of the visual system, such as retinotopic projections, is a distinguished feature of the sensory nervous system. However, despite numerous relevant studies, several fundamental questions remain to be answered. For instance, although the retinotopic organizations of the dorsal lateral geniculate nucleus (LGd) and the primary visual cortex (VISp) have been reported in separate studies (ref. 35 for LGd; ref.34 for VISp), no observation has yet provided a clear description of the spatial organization of the feed-forward projections from the LGd to the VISp at the system level. Furthermore, although the visual topography is maintained in the feedforward projections from VISp to the extrastriate visual areas in mice ^36, 37^, whether such topographic organization is maintained in the feedback connections from the higher cortical areas to VISp remains to be answered **(Fig. 4a)**. Therefore, AMaSiNe was used to observe spatial organization of the visual system in mice.

**Figure 4.**
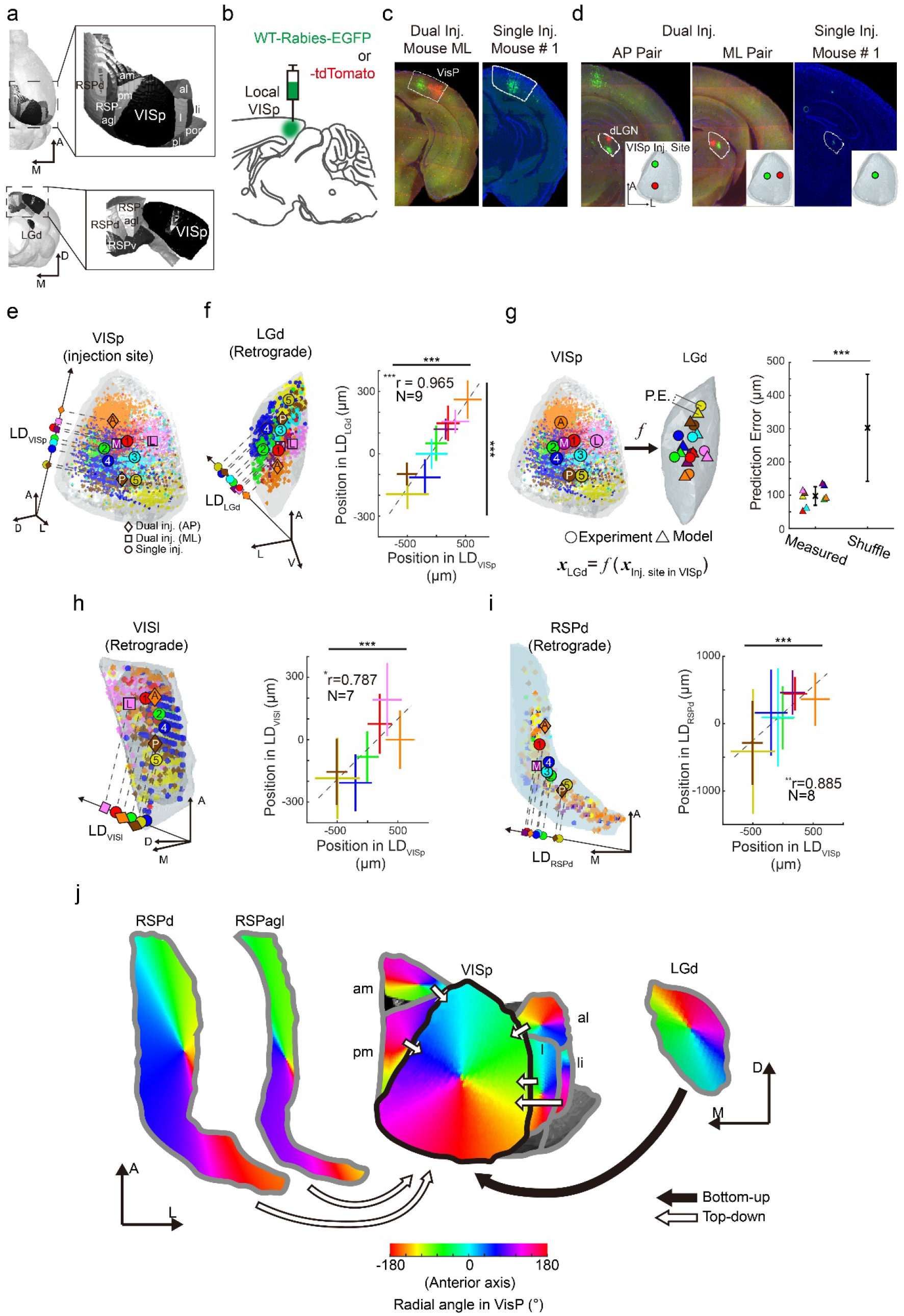
Topographical organization of neural projections to VISp. (a) Visual regions projecting to VISp. (b) Strategy for WT-Rabies injection for retrograde tracing of the inputs to VISp. (c) Injection sites made in local VISp regions. (d) Retinotopic organization of traced neurons in LGd. (e) Distributions of labeled neurons in VISp obtained from 9 injection sites (5 mice). Data obtained from multiple mice are drawn on the common reference space (see Supplementary Fig. 3). The center-of-mass positions of labeled neurons in each data set is projected onto the linear discriminant vector (LD). (f) Retinotopic projection from LGd to VISp. The distributions of labeled neurons in LGd are reconstructed in 3-D (left). Note that labeled neurons were spatially well segregated in both regions (right; mean ± SD; one-way ANOVA, ***p<0.001). The relationship between the neuronal distribution in VISp and in LGd shows strong correlation, indicating robust retinotopic connection from LGd to VISp (Pearson’s correlation analysis, mean ± SD; ***p<0.001). (g) Linear regression model of the spatial relationship of the feedforward projection from LGd to VISp. (left). The accuracy of model was tested by comparing the prediction error (P.E.) between observed data and randomly shuffled control (right; mean ± SD; Wilcoxon rank sum test, ***p<0.001, n = 9 and 500 for model and shuffled, respectively). (h and i) Spatial organization of feedback to VISp from VISl and RSPd (mean ± SD; ***p<0.001 for one-way ANOVA; *p<0.05, **p<0.01 for Pearson’s correlation analysis). (j) Summarized topography of input projections to VISp. Spatial arrangement of bottom-up (filled arrow; from LGd) and top-down (hollow arrow; from VISam, VISpm, VISal, VISli, RSPd, and RSPagl) projections to VISp was quantitatively estimated and visualized based on the results shown in Figures 4e–i (see Supplementary Fig. S4 and S5 for input from extrastriate and retrosplenial areas).

We injected wild-type rabies virus (RV) expressing either EGFP or tdTomato in local regions of mouse VISp to retrogradely trace the source of long-range inputs to VISp (**Fig. 4b**; 200 nL per injection; single injection of RV-EGFP in 5 mice; AP- or ML-paired dual injection of RV-EGFP and -tdTomato in 2 mice). Imaged data sets collected from individual RV-injected brains were analyzed with AMaSiNe. Analysis results showed that all injections were successfully targeted at local VISp regions without any spillovers into neighboring regions and long-range inputs were successfully traced **(Fig. 4c–d)**.

In the previous section, we showed the 3-D reconstructed brain accurately fit into ARA. This indicates that data from different mouse brains can be merged into a common reference space. Therefore, we directly compared the neuronal distribution obtained from multiple brains in ARA (**Fig. S3** shows further justification for this analysis). We first extracted the linear discriminant vector (LD) indicating the direction where the 3-D distributions of labeled neurons from multiple mice are best segregated 38 and projected detected neurons onto the LD in each ROI (**Fig. 4e, f, h, and i** for VISp, LGd, lateral area in the extrastriate cortex (VISl), and dorsal region of the retrosplenial cortical area (RSPd), respectively). Then, the spatial organization of neuronal distribution in VISp and in each of its input sources was investigated by examining the center-of-mass positions of labeled neurons in each data set in VISp and in other ROIs (**Fig. 4f, h and i** for VISp-LGd, VISp-VISl, VISp-RSPd relationships, respectively; ***p<0.001, **p<0.01, *p<0.05, Pearson’s correlation test). Results showed the distribution of labeled neurons in LGd, VISl, and RSPd was highly correlated with the distribution in VISp (r = 0.965, 0.787, and 0.885, respectively), indicating the topography of neural projections was robustly maintained in these connections. In addition, neurons providing inputs to distinct local VISp regions were spatially well segregated in all these regions, providing further evidence of systematic organization of the visual pathway (***p<0.001, one-way ANOVA). Among the 11 regions studied that projected to VISp (**Fig. 4a**), eight regions, except the posterolateral (VISpl) and postrhinal areas (VISpor) in the extrastriate cortex and the ventral part of the retrosplenial areas (RSPv), provided topographic local projections to VISp (**Fig. S4 and S5**).

Having observed an orderly projection from ROIs to VISp, whether the 3-D spatial relationship between VISp and its input sources can be explained with a simple model was investigated. We established a linear regression model that describes the relationship between the neuronal distribution in VISp and in each ROI (e.g. **Fig. 4g**, left for LGd). The center-of-mass positions of neurons in each ROI were used to model the relationship with minimum-norm solution. The accuracy of these models was validated by comparing the model prediction error with the distance of the random parings of the prediction and experimentally obtained data (**Fig. 4g**, right; Wilcoxon rank sum test, ***p<0.001, n = 9 and 500 for model and shuffle, respectively; **S4 and S5** for other ROIs). The spatial organization of the projection from 8 ROIs to the VISp was simulated using these models (**Fig. 4j**). Based on the results, the systematic organization of connections between major visual areas and VISp were quantitatively delineated. Compared to reports in previous studies showing retinotopic feedforward projections of VISp neurons to extrastriate cortical areas ^36, 39–41^, feedback projections from the higher visual areas were also spatially organized in a similar retinotopy, indicating the presence of reciprocal neural projections between local regions in VISp and those in extrastriate cortices.

## Discussion

To date, analysis of the mouse brain slice has remained subjective and highly susceptible to human error, leading to erroneous and inconsistent analysis results. In addition, the conventional analysis method is not only laborious but also inappropriate in practicality for direct comparison of data sets from different animals. Previous studies have taken various approaches to address these issues ^21, 22^, but the major problems of the conventional analysis with manual alignment and calibration have not been yet resolved.

To address the major issues of mouse brain analysis, we developed AMaSiNe, a novel software for automatic annotation of labeled neurons in the mouse brain image. AMaSiNe automatically locates the precise positions of experimentally obtained images in the common reference space, ARA, records the locations of labeled single neurons and calibrates data sets obtained from different animals on a common 3-D space for direct comparison. Unlike previous algorithms that match different mouse brains on a volume-to-volume basis for comparison ^21, 42^, AMaSiNe operates on an image-to-image basis, saving time and effort for preparing whole-brain slice images. Due to the image-to-image comparison algorithm, our software finds accurate alignment parameters of slices, including yaw and pitch angles of sectioning and AP positions of individual sections, with only a couple of single slices allowing the 3-D reconstruction of the labeled neurons onto a brain that precisely fits into ARA, a standard atlas. Furthermore, the algorithm is versatile in fitting images of brain slices stained using various methods, such as Nissl, DAPI, and autofluorescence. Therefore, investigators have more options for choosing staining methods optimized for their experimental purposes. Finally, the software was currently built with MATLAB, but users may easily adjust, convert, or add subjections as needed (e.g. combining with a deep neural network for detailed neurite detections) ^32^. In addition to customizing the algorithm, importing other versions of reference atlases for the analysis of brain images from other species is possible. Overall, AMaSiNe provides a powerful analysis framework for the automated mapping of neural circuits in a standard space with minimum effort.

To evaluate the performance of AMaSiNe, we used its accuracy to quantitatively determine the spatial organization of neural projections in the visual system, including projections from LGd, extrastriate areas, and retrosplenial cortices to VISp. By estimating spatial organization of inputs from multiple regions converging to a common target region of VISp, AMaSiNe showed a systematic analysis of delicate neural wirings in the mouse visual system. Consequently, topographic organization of seven extrastriate areas were identified with five providing region-specific inputs to the VISp. The spatial arrangements of these feedback connections were similar to the feedforward projections from VISp ^36,39–41,43^, showing the presence of reciprocal loops between visual areas in the cortex. Although the retinotopic organization in LGd 35 and the spatially organized feedback to VISp from the lateral visual area in the extrastriate cortex ^44^ have been previously reported, to the best of our knowledge, this is the first precise observation of the topographic structures of the projections to VISp throughout the whole visual areas, which is crucial to investigate how neurons communicate to perform appropriate functions in the visual system ^43, 45, 46^.

Our results show that AMaSiNE allows a complicated spatial organization of bottom-up and top-down connections of neural circuits to be found in a 3-D brain, implying that more in-depth insights for understanding the function of brain circuits can be achieved by facilitating accurate and quantitative analysis of the spatial organization of neural circuits in any region of the brain.

## Methods

### Virus Production

For retrograde tracing, we produced a rabies virus coated with G protein and containing either a red fluorescent protein (RV-∆G-tdTomato) or a green fluorescent protein (RV-∆G-eGFP) instead of G protein encoding gene in B7GG cells based on the previously published protocol by Osakada and Callaway^47^. To describe the production procedures briefly, B7GG cells were infected with the seed virus (RV-∆G-tdTomato or RV-∆G-eGFP; day 0). On day 4, supernatant (1^st^ harvest) including viral particles coated with G protein was harvested and fresh media added. Supernatant was also collected on day 7 (2^nd^ harvest) and day 10 (3^rd^ harvest) using the same protocol. Next, all the viral supernatants (1^st^, 2^nd^, and 3^rd^ harvest) were harvested using 0.20 μm pore size filter (Whatman). The filtered virus supernatant was concentrated using ultracentrifugation at 19,200 rpm for 2 hours (Beckman, SW32 Ti rotor), media was removed from the pellet with viral particles, and the pellet resuspended with autoclaved PBS (LPS solution, #CBP007A). HEK 293T cells were infected with the concentrated virus to quantify the virus titer (8.1 × 10^9^IU/mL for RV-∆G-tdTomato, 1.2 × 10^10^ IU/mL for RV-∆G-eGFP).

### Surgical Procedures and Animal Handling

The care and experimental manipulation were conducted in accordance with KAIST Institutional Animal Care and Use Committee (IACUC-14-145). Adult female WT mice (C57BL/6J, #000664, P56–P70) were anesthetized *via* inhalation of isoflurane (3% induction and approximately 1.5 – 2% maintenance) (Terrel) and head-fixed to a stereotaxic frame. Throughout surgery, body temperature was maintained at 37°C and eyes were covered with Vaseline to prevent drying. A midline incision was made on the scalp and tissue was cleared from the cranium. Bregma and lambda fissures were identified under the stereoscope and the mouse head was adjusted until the difference between the dorsoventral coordinates of bregma and lambda was less than 10 μm. For each mouse, one (single tracing with RV-∆G-eGFP only; 5 mice) or two (dual tracing with both RV-ΔG-eGFP and RV-ΔG-tdTomato) small craniotomies were performed above local regions of the right VISp; each mouse was assigned one or two of the four different injection target sites in local VISp: anterior site coordinates (AP, ML from bregma, and DV from pia surface; −3.1 mm, −2.6 mm, and 0.4 mm, respectively), posterior site coordinates (−4.3 mm, −2.6 mm, and 0.4 mm, respectively), medial site coordinates (−3.7 mm, −2.2 mm, and 0.4 mm, respectively), lateral site coordinates (−3.7 mm, −3.0 mm, and 0.4 mm, respectively).

At each target site, approximately 200 – 250 nL of either RV-ΔG-eGFP or RV-∆G-tdTomato was injected at the speed of 18.4 nL/min. After each injection, injection pipettes were left undisturbed for 10 – 15 min for sufficient absorption of the viral solution by the tissue. Next, pipettes were slowly removed and the scalp was sutured with glue (3M, #1469SB). Postoperative analgesia was provided *via* EMLA (2.5% lidocaine/prilocaine) application to the sutured skin. Mice were placed in a heated box under human observation until fully awake from anesthesia and then single-caged.

### Tissue Preparation

Five days after the RV injection, mice were overdosed with isoflurane and perfused *via* the ascending aorta with 15 mL of 0.1 M PBS (pH 7.4; LPS solution, #CBP007A) solution followed by 25 mL of 4% w/v paraformaldehyde (PFA, Sigma-Aldrich #158127) in 0.1 M PBS. Brains were collected and left in 4% w/v PFA solution for 4 hours at 4°C. After post-fixation, the brains were placed in 5 mL of 30% sucrose solution (wt/vol) (Sigma Aldrich, #S9378) and placed on a shaker at room temperature (RT) for 1 – 2 days. Next, the brains were dehydrated and embedded in optimal cutting temperature (OCT) medium (Tissue-Tek #4583) and frozen at −80°C. Embedded brains were cryosectioned (Leica, CM1520) and 40 μm-thick tissues were collected. We ensured that coronal sections containing at least LGd, RSP, and visual cortical areas were all collected for imaging. Collected tissues were washed three times for 10 min with phosphate buffer (PB, 0.1M) and mounted using mounting medium with DAPI (Vector Labs, #H-1200).

For the verification of AMaSiNe when analyzing tissues stained with Nissl, brain tissues were collected from a single brain (approximate AP range from −6 – 3 mm, which covers most of the cortical regions and part of the cerebellum) that was not injected with any tracer or virus. Tissues were cryosectioned as described above. Collected tissues were washed three times for 15 min in 0.1M PBS and permeabilized with 0.1% Triton X-100 (Sigma Aldrich, #X100) in 0.1 M PBS for 10 min. Then, the tissues were washed twice for 5 min with 0.1 M PBS and treated with 200 μL of diluted NeuroTrace (Thermo Fisher, #N21480) in 0.1 M PBS (1:100). Treated tissues were incubated for 20 min at RT. Following incubation, the tissues were washed with 0.1% Triton X-100 in 0.1 M PBS for 10 min. After additional two 10-min washings and a single two-hour washout with 0.1 M PBS at RT, tissues were mounted with mounting medium containing DAKO.

### Imaging

Fluorescent images of prepared samples were obtained using a slide scanner microscope (10x; 0.653 μm pixel resolution; 0.45 NA; Zeiss, AxioScan.Z1). When imaging, proper excitation wavelengths were used for different fluorophores (Zeiss, HXP120V): 358 nm for DAPI, 488 nm for eGFP, and 554 nm for tdTomato and autofluorescence. Image files were exported in 8-bit .tif format after contrast normalization and downsampling to 1.306 μm pixel resolution.

### Development of AMaSiNe

AMaSiNe was built using MATLAB R2017a with Computer Vision System Toolbox (R2017a release), Image Processing Toolbox (R2017a release), and Parallel Computing Toolbox (R2017a release) on Windows 10. The 25 μm resolution Allen Mouse Common Coordinate Framework (CCFv3; for angle searching and image registration of autofluorescent images) with the Nissl volume (for angle searching and image registration of DAPI- or Nissl-stained images), and structural annotation volume were downloaded from the Allen Brain Atlas data portal (http://brain-map.org).

### Image Preprocessing and Position-Parameter Searching

Exported brain images in *.tif format went through a series of preprocessing stages. Original images were first binarized and only the largest object in the image, namely the brain slice, was left to minimize the influence of noise or dirt outside the brain slice for further processing.

For the slicing-angle searching stage, images were sampled at approximately 250 μm – 1,000 μm interval from the whole series of slice images from a single brain (anchor images). This measure was performed to reduce the computational time. Sampled images were further preprocessed: top and bottom 1% of all pixel values were saturated to normalize the intensity of brain slice image and the intensity-adjusted image was sharpened with an unsharpened mask.
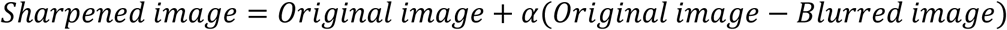

where α is the amount of sharpening (α = 5) and the blurred image is the original image filtered with a Gaussian mask (σ = 75 μm; filter size = 400 μm × 400 μm). Finally, sharpened images and ARA volume were downsampled to the 50 μm pixel (voxel) resolution with bicubic interpolation to further reduce computational time; the ARA volume was downsampled only for this stage to find the position parameters of individual brain slices.

Before a preprocessed anchor image and an ARA slice were compared, the preprocessed anchor image was structurally adjusted twice to roughly match its shape to the ARA slice for comparison. First, SURF points (number of octaves = 1) ^27^ were extracted from both images and extracted feature points were described with HOG descriptors (250 μm × 250 μm cell size, 13 × 13 blocks, 7 × 7 cell overlap between adjacent blocks) ^28^. Feature points with similar HOG descriptor vectors in both images were then matched. A transformation matrix for non-reflective similarity transformation was computed with the matched feature points in both images. Using this matrix, the preprocessed anchor image was registered linearly. The registered image then went through a second transformation with the same method described previously but with an affine transformation.

For the actual computation of the ISS, grid points were sampled with 200 μm intervals and SURF points were extracted from anchor and ARA slice images, respectively. Feature points in both images were described with HOG descriptors (200 μm × 200 μm cell size, 21 × 21 blocks, 11 × 11 cell overlap between adjacent blocks) and similar feature points were matched. ISS was then computed with the HOG descriptors of the matched feature points in both images (*HOG_matched feature in obtained image_* and *HOG_matched feature in ARA slice_*) based on the following equation:

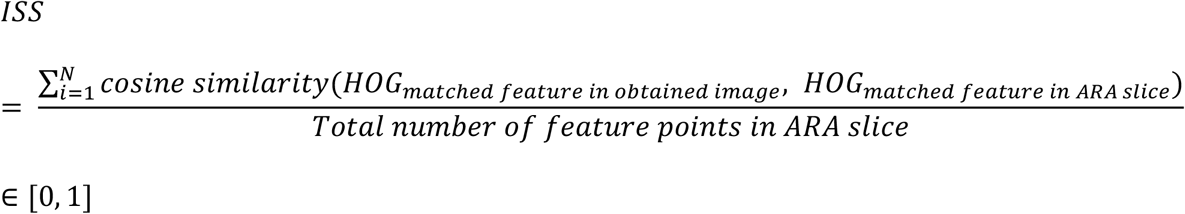

where N is the total number of matched features in both images and cosine similarity of two HOG descriptor vectors is defined as follows:

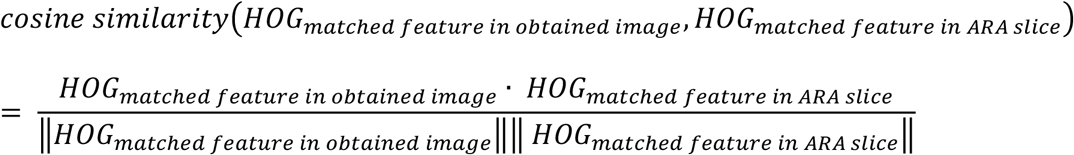

Prior to searching the position parameters of individual anchor images with the three-step search algorithm, the search-starting position of each image was set as follows: 3-D ARA was sliced at an interval of 250 μm at both the slicing yaw and tilt angle values of 0**°**. The resulting 52 ARA slice images were compared to the obtained anchor image as described above, and the AP position of the ARA slice with the highest ISS was set as the starting point for further position searching.

Once the positions of the anchor images were found, the AP positions of non-anchor images (images not used for the slicing angle searching) were interpolated between the nearest two anchor images at equal distances. The slicing yaw and pitch angles for non-anchor images were set as the values found with the anchor image because the individual brains had been firmly mounted on the sectioning apparatus and not repositioned during sectioning.

### Image Warping and Cell Detection

Each obtained image was structurally transformed to match the shape of the ARA image sliced at the corresponding position. Obtained images were first downsampled to the 25 μm resolution to match the ARA slice image. Features from both images were extracted and matched in a similar manner to the one used in the position searching step. Three transformation matrices, one non-reflective similarity transformation followed by two affine transformations, were computed and temporarily saved for image registration of an obtained image at its original resolution. The positions of matched features of the ARA slice and obtained image that had been linearly transformed three times were also temporarily saved to be used as control points during non-linear transformation at original resolution. The transformation matrices and feature points computed at 25 μm resolution were rescaled to the original resolution of the obtained image and applied. For non-linear transformation, the local-weighted-mean registration algorithm ^29^ was used.

Neurons labeled with tdTomato or eGFP in warped images were located as follows: First, the radius range of the soma to be detected was set as approximately 8 – 16 μm. Then, a Fermi filter, F_Fermi_, was applied to remove salt-and-pepper noises

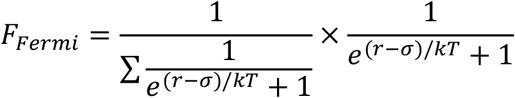

where *r* is set as the radius of the 50 μm × 50 μm mask size (70 μm), *σ* value set as the lower boundary of the soma size (8 μm), and *kT* set as 5. Next, a difference-of-Gaussian filter, F_DoG_ was applied to boost soma-like structures in the image:

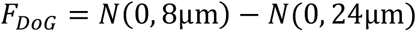

where N(μ, *σ*) is a Gaussian distribution with mean μ and standard deviation *σ*. Finally, labeled somas were located using the two-stage method after Hough transformation of the preprocessed image ^30, 31^. Detected neurons were marked with red dots on warped images and visually inspected. Undetected neurons and false-positively detected neurons, though not many (data not shown), were manually added or deleted with a graphic user interface built in MATLAB. The locations of detected somas in each image were saved as a separate .mat file for annotation.

### 3-D Reconstruction and Cell Annotation

Registered images and the positions of detected neurons in each image were first downscaled to 25 μm to match the resolution of ARA volume and aligned along the slicing axis. Because the resulting 3-D reconstructed brain does not precisely fit into the ARA volume due to the tilted slicing of the brain during tissue preparation, the 3-D positions of neurons detected were compensated for the position deviation from ARA.

***Position***_*adjusted*_ **= *M***_*yaw*_***M***_*pitch*_ × (***Position***_*unadjusted*_ – ***pivot***) **+ *pivot*** where ***Position*** _*adjusted*_ and ***Position***_*unadjusted*_ are the 3-D coordinates of neurons before and after tilt-angle compensation, respectively, ***Pivot***, the center coordinate of ARA volume, and ***M***_*yaw*_ and ***M***_*pitch*_ the 3 × 3 matrices to compensate the tilt in yaw and pitch.
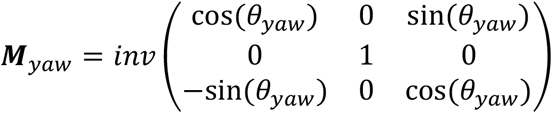

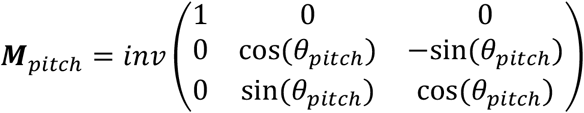

To describe the position of neurons in the commonly used AP-ML-DV coordinate frame, we set the 214^th^(x)-348^th^(y)-116^th^ (z) pixel of the ARA volume as the bregma coordinate; the adjusted positions of neurons were translated by [-348, 116, 214] pixels and upscaled to its original resolution.

For the annotation of located neurons, individual pixel coordinates of the ARA annotation volume were translated using the bregma coordinate and multiplied by 25 μm. Then, the distance between each adjusted position of detected neurons and all ARA annotation pixels were measured. The ARA coordinate with the smallest distance was set as the ROI the labeled neuron belonged to; the located neurons were annotated accordingly.

### Analysis of the Spatial Organization of Inputs to ViSp

Using the 3-D location data of labeled neurons annotated (9 data sets from 5 mice), we studied the relationships between the spatial organization of neurons in VISp and in each ROI projecting to VISp. To ensure statistical robustness of analysis, ROIs in each data set that contained less than 10 labeled neurons were excluded. From the valid data sets, a multi-class linear discriminant vector (LD)^38^ in each ROI were computed. Neurons in each ROI were projected onto the LD to produce a 1-D distribution on LD. The center-of-mass (i.e. the mean position) of LD-projected neurons in each ROI of each data set was calculated. To study the relationships between the distribution of neurons in VISp and in ROIs projecting to VISp, the Pearson’s correlation coefficient with the center-of-mass position was measured in all valid data sets in each region.

The 3-D linear regression models for the spatial relationship between VISp and its input sources were built with minimum-norm solution using the center-of-mass position of each data set in each ROI. The model errors were computed by measuring the distances between the prediction results and their corresponding center-of-mass positions obtained from actual data sets. The accuracy of the models was validated by comparing the model prediction errors with the distance of the random parings of the prediction and experimentally obtained data.

The overview of the spatial organization of inputs to VISp (**Fig. 4j**) were designed as follows: The point coordinates spaced regularly at 25 μm intervals in layers II to IV of VISp in ARA were used as inputs to the model equations. Because the models were linear, numerous output points deviated from the outer boundaries of corresponding ROIs (e.g. RSPd and RSPagl); such points were excluded when plotting the results. In addition, several parts of ROIs were not fully covered by the model outputs, thus, the existing output points were linearly extrapolated to fully cover each ROI.

## Acknowledgements

We thank Yang Dan and Kwanghun Chung for providing helpful comments in the early stages of this study. This research was supported by the Basic Science Research Program through the National Research Foundation of Korea (NRF) funded by the Ministry of Science and ICT (NRF-2016R1C1B2016039, NRF-2016R1E1A2A01939949 to S.P. and NRF-2017R1A2B3008270 to S.L.) and the High Risk High Return Project of KAIST (N11170092 to S.P.).

## Author contributions

J.S. designed the project, developed software for analysis, performed the animal surgeries and imaging experiments, analyzed data, and wrote the manuscript. Y.S. performed the animal surgeries and imaging experiments, analyzed data, and wrote the manuscript. J.K. performed the animal surgeries and imaging experiments and wrote the manuscript. W.C. analyzed data and edited the manuscript. S.L. designed the project, directed the experimental research, and edited the manuscript. S.P. designed the project, directed the software development and analysis research, and wrote the manuscript. All authors discussed and commented on the manuscript.

## Competing interest declaration

The authors declare no competing interests.

